# Utilizing Digital Pathology to Quantify Stromal Caveolin-1 Expression in Malignant and Benign Ovarian Tumors: Associations with Clinicopathological Parameters and Clinical Outcomes

**DOI:** 10.1101/2021.08.18.456818

**Authors:** Daryoush Saeed-Vafa, Douglas C. Marchion, Susan M McCarthy, Ardeshir Hakam, Alexis Lopez, Robert M. Wenham, Sachin M. Apte, Dung-Tsa Chen, Anthony M. Magliocco, Jonathan M. Lancaster, Brett M. Reid, Jennifer B. Permuth

## Abstract

Loss of stromal caveolin-1 (Cav-1) is a biomarker of a cancer-associated fibroblast (CAF) phenotype and is related to progression, metastasis, and poor outcomes in several cancers. The objective of this study was to evaluate the clinical significance of Cav-1 expression in invasive epithelial ovarian cancer (OvCa). Epithelial and stromal Cav-1 expression were quantified in serous OvCa and benign ovarian tissue in two, independent cohorts – one quantified expression using immunohistochemistry (IHC) and the other using multiplex immunofluorescence (IF) with digital image analysis designed to target CAF-specific expression. Cav-1 expression was significantly downregulated in OvCa stroma compared to non-neoplastic stroma using both the IHC (p=0.002) and IF (p=1.8×10^-13^) assays. OvCa stroma showed Cav-1 downregulation compared to tumor epithelium with IHC (p=1.2×10^-24^). Conversely, Cav-1 expression was higher in OvCa stroma compared to tumor epithelium with IF (p=0.002). There was moderate correlation between IHC and IF methods for stromal Cav-1 expression (*r*^2^= 0.69, p=0.006) whereas there was no correlation for epithelial expression (*r*^2^=0.006, p=0.98). Irrespective of the staining assay, neither response to therapy or overall survival correlated with the expression level of Cav-1 in the stroma or tumor epithelium. Our findings demonstrate a loss of stromal Cav-1 expression in ovarian serous carcinomas. Studies are needed to replicate these findings and explore therapeutic implications, particularly for immunotherapy response.

## Introduction

Ovarian carcinoma (OvCa) is the fifth leading cause of cancer-related deaths among women in the United States. In 2020, an estimated 21,750 new cases of OvCa and 13,940 related deaths occurred, making it the most deadly of all gynecologic cancers [1]. Due to non-specific symptoms and lack of early detection methods, nearly 80% of women present with advanced stage (III/IV) disease with a 5-year relative survival rate of approximately 30% [1]. Despite standard treatment, which includes surgical debulking followed by platinum-based chemotherapy, most women succumb to persistent or recurrent platinum-resistant disease. Thus, there is a dire need to discover novel biomarkers for OvCa that are associated with clinical outcomes and could potentially serve as therapeutic targets.

The tumor microenvironment is a critical component of ovarian carcinogenesis and immune response that can yield information relevant to prognosis and chemoresistance [2-7]. Cancer-associated fibroblasts (CAFs) are the most abundant cells in the tumor microenvironment and are known to promote tumor growth, invasion, and metastasis [8-12] and impact the efficacy of chemotherapy [8, 10-13]. Ovarian CAFs are genetically more stable than tumor cells [14], making them less prone to developing chemoresistance and therefore excellent therapeutic targets. However, the clinical significance of proteins associated with CAFs in OvCa is largely unexplored.

Caveolin-1 (Cav-1) belongs to a family of scaffolding proteins that regulate signal transduction and cell proliferation [15] and has emerged as a marker of a lethal CAF phenotype. Specifically, the absence or loss of stromal Cav-1 expression has been shown to activate epithelial AKT signaling [13, 16] and be a powerful predictor of poor outcomes in numerous malignancies, including breast, prostate, pancreatic, papillary thyroid cancers, and melanoma [8-15, 17-20]. Furthermore, the reduced expression of Cav-1 in the ‘reactive’ or desmoplastic stroma of malignant tumors correlates with more aggressive disease and advanced stage [12-14]. Investigations in OvCa have found Cav-1 functions as a tumor suppressor gene and is significantly downregulated in ovarian cancer cell lines and carcinomas which promotes tumor growth, survival, invasion, and metastasis [21-25]. Additionally, stromal Cav-1 expression has been detected in benign ovarian serous tumors but is largely absent in malignant and borderline ovarian serous tumors [26]. Several studies have observed loss of Cav-1 staining specifically in OvCa stromal cells, suggesting Cav-1 may be a marker of CAFs and poor survival in OvCa, as in other cancers. Supporting this role, reduced Cav-1 mRNA expression has been associated with poor survival in serous carcinomas from The Cancer Genome Atlas. Further, stromal Cav-1 expression has been correlated with more aggressive ovarian serous tumors [6, 26]. However, a recent study evaluating Cav-1 expression in OvCa stroma did not detect an association with survival [26] but this may be due to the small sample size (n=31) or quantification of expression as absent or present. Thus, it is unclear whether stromal Cav-1 expression could serve as a prognostic and therapeutic marker for OvCa.

The purpose of this study was to comprehensively quantify stroma and epithelium Cav-1 expression and to explore the relationship with clinicopathologic variables, response to first-line platinum-based chemotherapy, and survival. We evaluated tissue microarrays (TMAs) comprised of human serous carcinomas and benign ovarian tissue from two independent, relatively large cohorts of women diagnosed and treated at our National Cancer Institute-designated Comprehensive Cancer Center. We evaluated one cohort using an immunohistochemistry (IHC) assay and semi-quantitative scoring and the other with a multiplex immunofluorescence (IF) assay quantified using automated digital image analysis because of its more precise and objective quantification of protein expression[27].

## Materials and Methods

### 2.1 Study populations and clinical data

The study population included two cohorts diagnosed and treated at Moffitt Cancer Center, with the entire project conducted in accordance with full institutional review board approval. This study has been carried out in accordance with the ethical principles for medical research involving human subjects, set by the Declaration of Helsinki and was approved by the University of South Florida Institutional Review Board (IRB#: Pro00008518). Each cohort included subjects with malignant and benign disease. Malignant cases were defined as women with primary pathologically-confirmed invasive epithelial OvCa treated with surgery followed by platinum-based therapy. Benign cases included women who underwent surgery for benign gynecologic conditions. Both groups of women donated tissue specimens under the auspices of an institutional tissue banking protocol. The first cohort (Cohort 1) comprised subjects who underwent surgery between 1987 and 2010, whereas the second cohort (Cohort 2) underwent surgery between 2000 and 2012. Since OvCa incidence and mortality is highest in non-hispanic whites^1^ and most OvCas are of the serous subtype, which is etiologically distinct from other histologies [28], our analyses focused on invasive serous adenocarcinomas among non-Hispanic whites.

Demographic and clinical data was obtained from the Moffitt Cancer Registry (age at diagnosis, histology, pathologic grade and stage, CA-125 levels, debulking status, treatment regimens and response, recurrence) supplemented by the medical record, and the National Death Index (for vital status). For purposes of this study, overall survival was defined as the time between the date of diagnosis and the date of death or last contact. Treatment response was retrospectively determined using standard criteria for patients with measurable disease using World Health Organization guidelines [28]. CA-125 was used to classify response in the absence of a measurable lesion [29, 30]. Complete clinical responders (CR) had complete disappearance of all measurable disease for 4 weeks after completing first-line platinum-based chemotherapy, or in the absence of measurable lesions, a normalization of the CA-125 level for at least 4 weeks after therapy and were therefore initially chemosensitive. Patients who demonstrated a partial response (defined as a 50% or greater reduction in tumor burden obtained from measurement of each bi-dimensional lesion for at least 4 weeks, or a drop in CA-125 by >50% for at least 4 weeks after completing therapy), no response (had stable disease), or progressed during adjuvant therapy were considered chemoresistant and thus classified as incomplete responders (IR).

### 2.2 Tissue microarrays

We utilized two ovarian tissue microarrays (TMAs) in this study. The first was an existing TMA (TMA 1) of Cohort 1 subjects that included 100 OvCa cases selected based on treatment response (50 CRs and 50 IRs) and 35 benign cases. The second (TMA 2) was constructed specifically for this investigation from Cohort 2 subjects that included 128 unselected OvCa cases and 30 benign cases. TMA 2 included a subset of OvCa cases (n=11) sampled in TMA 1/Cohort 1. Cores for both TMAs were obtained from formalin-fixed paraffin-embedded tissue blocks. Hematoxylin-and-eosin (H&E) stained slides from selected blocks were reviewed by a pathologist with expertise in gynecologic tumors (AH, AL) to confirm the diagnosis, in accordance with the 2018 WHO guidelines, and select regions of interest (ROI). For the established TMA 1, OvCa sampling included single cores punched from macrodissected tumor tissue. Benign case samples included single cores from normal ovarian tissue. For TMA 2, four ROIs were preferentially sampled for OvCa cases: 1) abundant CAF-enriched tumor stromal cells immediately adjacent to the tumor epithelium; 2) areas of abundant tumor epithelial cells; 3) paired adjacent normal ovarian and/or fallopian epithelial tissue; 4) normal non-neoplastic stroma with a high abundance of normal fibroblasts that are morphologically distinct from CAFs [8, 9]. For each ROI on TMA 2, duplicate 1.0 mm cores were sampled to account for tissue heterogeneity and the cores were placed side-by-side on a recipient microarray block. Samples for benign cases in TMA 2 included multiple cores from normal stroma and/or epithelium tissues, when present. The Moffitt Tissue Core Facility constructed the TMAs using a precision instrument (Tissue Arrayer, Beecher Instruments, WI) per standard protocols.

### 2.3 Immunohistochemistry and scoring, TMA 1/Cohort 1

A rabbit polyclonal anti-Cav-1 IgG antibody (Santa Cruz Biotechnology, Santa Cruz, CA) was used for immunohistochemical (IHC) staining of 3 micron thick sections of the TMA block as previously described [31]. A board-certified pathologist (AH) scored the stains semi-quantitatively by multiplying the mean signal intensity scores by the percentage of positive cells. ROIs were identified and scores were calculated separately for each ROI. Tumor samples were scored for tumor epithelium and adjacent stroma. Benign samples were scored for normal stroma and epithelium regions. Staining intensity was scored as follows: 0 (negative; no staining), 1 (weak), 2 (moderate), and 3 (strong). The percentage of positive cells was scored as: 0 (0%), 1 (1-33%), 2 (34-66%), and 3 (67-100%).

### 2.4 Immunofluorescence and Digital Image Analysis, TMA 2/Cohort 2

A rabbit polyclonal anti-Cav-1 IgG antibody (Santa Cruz lot: G0314) was used for immunofluorescent (IF) staining of TMA 2 via a standard multiplexing protocol (S1). The AQUA^®^ system was used to perform digital image analysis (DIA) on the stained virtual slides generated by the Aperio ScanScope^®^FL fluorescent scanner. We designed an algorithm to target and quantify Cav-1 expression within the fibroblasts (i.e. the stroma compartment) and the tumor epithelial cells (i.e. the tumor epithelial compartment). The algorithm was designed such that the stroma compartment was defined by the presence of the mesenchymal vimentin signal (fibroblast intermediate filament stain; local to cytoplasm) and the DAPI signal (dsDNA stain; local to nucleus). However, vimentin is a ubiquitous mesenchymal marker that has the potential to stain a variety of cell types including tumor epithelial cells. To ensure that only the fibroblasts were considered in the Cav-1 stroma expression, pixels expressing both vimentin and pancytokeratin (epithelial stain; local to cytoplasm) were excluded from stroma compartment calculations. Ultimately, Cav-1 AQUA^®^ scores (nuclear, cytoplasmic, and complete cell (overall)) were obtained for the stroma compartment, as well as for the tumor epithelial compartment for each TMA core. The AQUA^®^ score was calculated from the sum of the target pixel intensity divided by the compartment area and normalized for image exposure time to make them directly proportional to protein concentration (molecules per unit area) [32]. For purposes of this study, only the overall scores (i.e. the total amount of Cav-1 expressed in the complete cell) within the different compartments (stroma or epithelium) were used for each core. For duplicate cores from the same participant, the mean compartment score was used for statistical analysis.

### 2.5 Statistical analysis

In our analysis, we excluded patients who received neo-adjuvant chemotherapy before surgery or had tissues sampled from recurrent disease. Further, cases diagnosed with histological subtypes other than serous were excluded (Cohort 1 n=3, Cohort 2 n=23). Cav-1 expression was compared by the following clinicopathologic features: age (young, ≤65 years, versus old > 65 years), pathologic grade (low-grade versus high-grade), pathologic stage (early-stage I/II versus advanced-stage III/IV), response to therapy (complete responders (CR)/ chemosensitive versus incomplete responders (IR)/ chemoresistant), and survival (long-term survivors of > 60 months versus short-term survivors of ≤ 36 months). For each cohort, the non-parametric Wilcoxon rank-sum test was used to compare the median difference in Cav-1 expression between clinicopathologic subgroups. Logistic regression was used to examine the association of Cav-1 expression with response to chemotherapy including adjustment for significant prognostic factors. The associations between Cav-1 expression and OS were visually assessed using Kaplan-Meier survival curves and log-rank tests. IHC score was dichotomized as the presence (1,2) or absence (0) of Cav-1, consistent with previous studies [31]. IF expression was dichotomized using the median expression as the cutoff. Cox proportional hazards regression was used to estimate the independent association of continuous Cav-1 expression with OS, controlling for significant clinical covariates.

## Results

### 3.1 Characteristics of study participants, Cohort 1

A total of 106 subjects were evaluated including 73 serous OvCa cases (Table 1) with chemo-naïve primary tumor tissue sampled and 33 controls with benign ovarian disease. All cases were treated with chemotherapy and 37 (51%) displayed a complete response, reflecting the TMA selection criteria. The average age at diagnosis was 62 (range: 33-83, median=63) and almost all were diagnosed at an advanced stage (III/IV, n=72) with high-grade (n=70) disease. Clinical follow-up for overall survival averaged 76.9 months after diagnosis (standard deviation, SD=91.4, range=3.6 to 392.9). A total of 65 (89%) of cases died during follow-up with 28 long-term and 36 short-term survivors.

**Table 1:** Immunohistochemical Cav-1 Expression by Clinical Variables in Cohort 1

| Outcome | N (%)<sup>a</sup> | Cav-1 Expression, median (SD) |  | Mean Δ (SD) |
| --- | --- | --- | --- | --- |
|  |  | Desmoplastic Stroma (DS) | Tumor Epithelium (TE) |  |
| Overall | 76 | 0.28 (0.73) | 2.50 (1.55) | -2.17 (1.31) |
| | | | | $p=4.3 \times 10^{-19}$ |
| Age at Diagnosis |  |  |  |  |
| ≤65 years | 43 (57%) | 0.26 (0.70) | 2.53 (1.45) | -2.19 (1.15) |
| >65 years | 33 (43%) | 0.30 (0.77) | 2.45 (1.70) | -2.15 (1.50) |
| | | $p = 0.82$ | $p = 0.94$ | $p = 0.85$ |
| Response to Therapy |  |  |  |  |
| Complete | 39 (51%) | 0.36 (0.74) | 2.44 (1.59) | -2.08 (1.38) |
| Incomplete | 37 (49%) | 0.19 (0.71) | 2.57 (1.54) | -2.28 (1.23) |
| | | $p = 0.08$ | $p = 0.73$ | $p = 0.33$ |
| Survival |  |  |  |  |
| Long-term (>60 months) | 30 (45%) | 0.37 (0.81) | 2.37 (1.71) | -2.00 (1.46) |
| Short-term (≤36 months) | 37 (55%) | 0.19 (0.71) | 2.57 (1.57) | -2.28 (1.28) |
| | | $p = 0.12$ | $p = 0.47$ | $p = 0.18$ |

<sup>a</sup> Survival groups do not total 76 due to cases with follow-up <36 months or death <60 and >36 months.

### 3.2 Expression of Cav-1 using immunohistochemistry, Cohort 1

Representative immunohistochemical staining of Cav-1 is shown in Figure 1. For both tumor and benign ovarian tissue, higher expression of Cav-1 was observed in the epithelial regions as compared to stromal regions. Tumor epithelial Cav-1 expression averaged 8-fold higher than adjacent stroma (mean 2.56 vs 0.29, p=8.7×10^-19^) while benign ovarian epithelium had 2.5 fold higher expression compared to normal stroma (mean 3.06 vs. 0.88, p=3.7×10^-9^). Expression of Cav-1 in benign stroma was two-fold higher than in tumor stroma (p=0.003). No difference in epithelial expression was observed between benign and tumor tissues (p=0.63).

**Figure 1.**
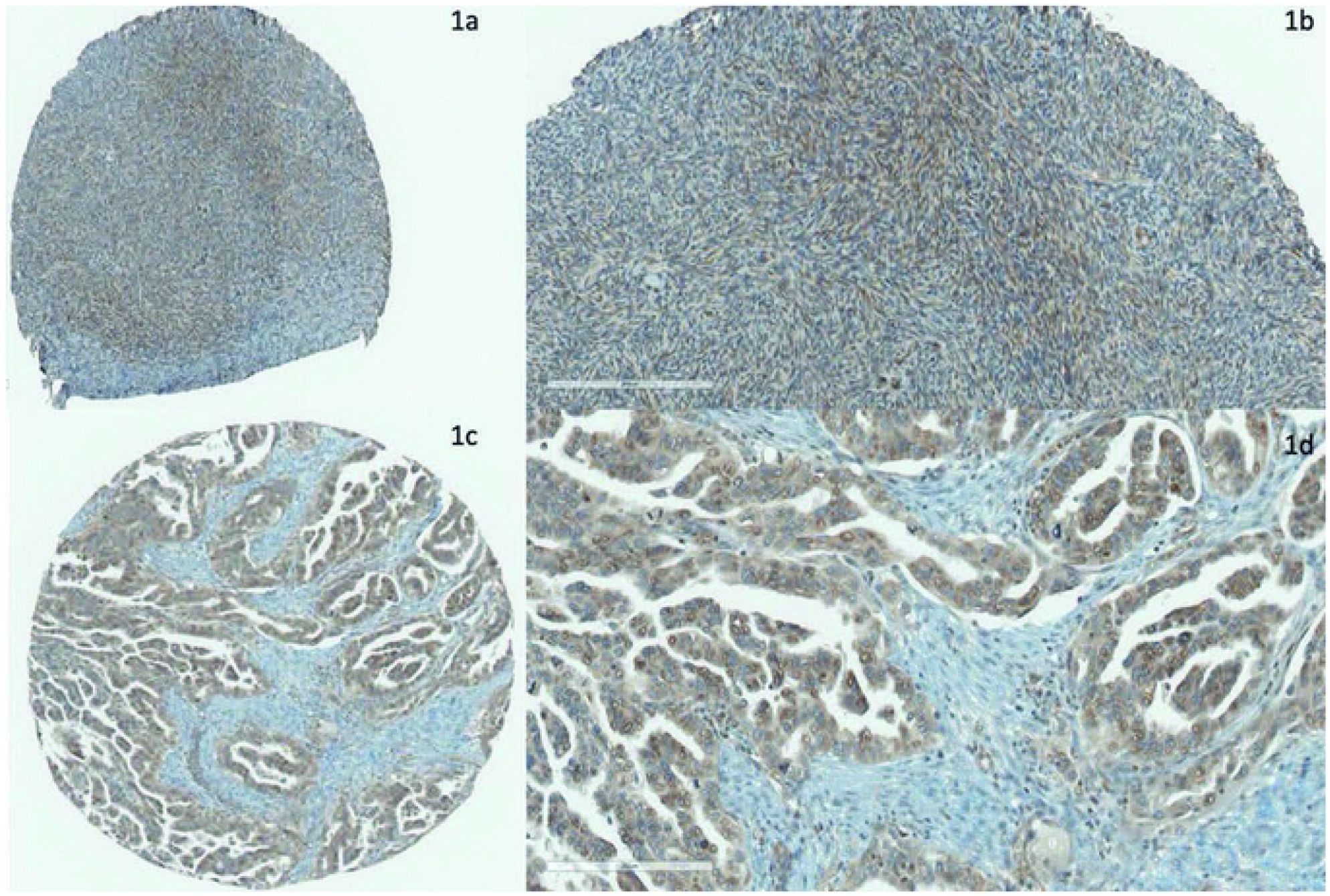
Immunohistochemical expression of Cav-1 in ovarian tissue. (A-B) Normal (benign) ovarian stroma stained with Cav-1 via immunohistochemistry. The fibroblasts in the benign ovarian stroma stain strongly positive for Cav-1. (C-D) High-grade ovarian serous adenocarcinoma stained with Cav-1 via immunohistochemistry. The tumor epithelial cells stain strongly positive for Cav-1, while the fibroblasts in the surrounding desmoplastic stroma stain weakly positive to negative for Cav-1.

Expression of Cav-1 was compared across clinicopathologic characteristics (Table 1). Cases with an incomplete response to therapy had a slightly lower expression of Cav-1 in the stroma as compared to those with a complete response (p=0.07). Additionally, short term survivors had a slightly lower expression of stromal Cav-1, but a slightly higher expression in the epithelium, as compared to long term survivors; however, these differences were not statistically significant. No difference was observed in the expression of Cav-1 in either the stroma or epithelium by age.

The median overall survival was 40.6 months for OvCa patients. For patients with presence of Cav-1 (Hscore>0) in the epithelium, survival time was comparable to those with an absence of Cav-1 (median survival 45.0 vs. 33.5 months, respectively, p=0.36; Figure 2a). Conversely, patients with presence of stromal Cav-1 expression had longer survival (median 73.3 months) as compared those with an absence of stromal Cav-1 (31.4 months), which was suggestive of an association (p=0.09, Figure 2b). However, after adjustment for age and response to therapy there was no difference in stromal Cav-1 expression by survival (Hazard Ratio, HR=0.83, 95% CI=0.58-1.19, p=0.31). Survival did not differ by epithelial Cav-1 expression in adjusted analysis (HR=0.99, 95% CI=0.84-1.16, p=0.89).

**Figure 2.**
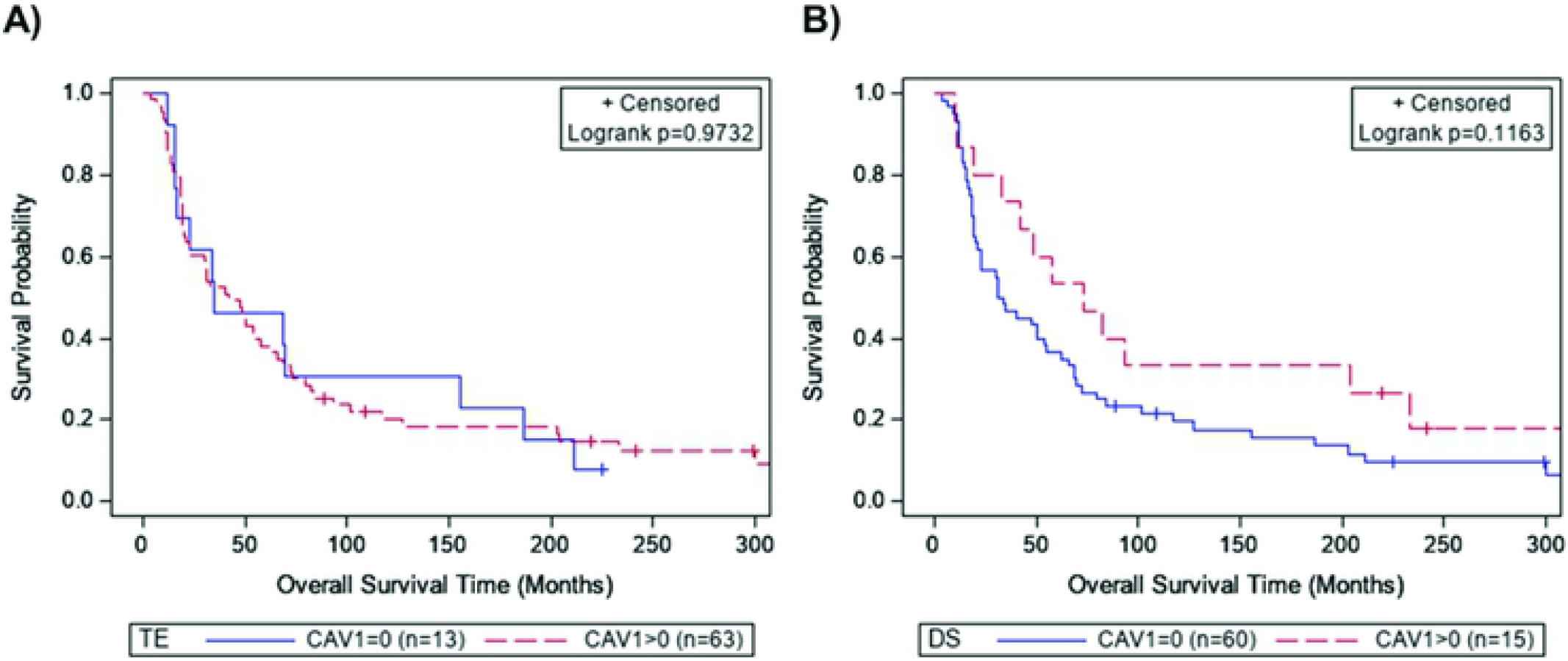
Overall survival by expression of Cav-1 in Cohort 1. Kaplan-Meier survival curves for Cohort 1 by presence of Cav-1 immunohistochemical expression in the (A) tumor epithelium and (B) stroma using an immunohistochemistry H-score of zero as a cutoff.

### 3.3 Characteristics of study participants, Cohort 2

A total of 135 subjects were evaluated including 105 with pathological diagnosis of serous OvCa (Table 2) and 30 controls with benign ovarian tissue. The average age at OvCa diagnosis was 60 (range: 24-83, median=62) and the majority presented with advanced (85%), high-grade (95%) disease, similar to our prior cohort. OvCa cases were unselected for response to therapy with most (77%) having a complete response. Additionally, 80 cases were optimally debulked. Patients were followed for an average of 60 months (SD=44.8, range=5.2 to 199.0) with 40 short-term and 45 long-term survivors.

**Table 2:** Immunofluorescent Cav-1 Expression by Clinical Variables in Cohort 2

| Outcome | N (%)<sup>a</sup> | Cav-1 Expression, median (SD) |  | Mean Δ (SD) |
| --- | --- | --- | --- | --- |
|  |  | Desmoplastic Stroma (DS) | Tumor Epithelium (TE) |  |
| Overall | 128 | 149.9 (44.5) | 124.4 (58.9) | 12.7 (45.9) |
|  |  |  |  | ***p* = 0.002** |
| Age at Diagnosis |  |  |  |  |
| ≤65 years | 78 (62%) | 140.0 (43.1) | 124.4 (54.1) | 10.2 (45.9) |
| >65 years | 48 (38%) | 154.0 (42.7) | 122.6 (66.3) | 15.5 (46.5) |
|  |  | ***p* = 0.01** | *p* = 0.29 | *p* = 0.53 |
| Grade |  |  |  |  |
| Low | 7 (5%) | 169.2 (58.5) | 184.2 (91.2) | -0.1 (62.1) |
| High | 121 (95%) | 149.3 (43.5) | 124.0 (56.1) | 18.7 (44.7) |
|  |  | *p* = 0.27 | *p* = 0.10 | *p* = 0.34 |
| Stage |  |  |  |  |
| Early (I/II) | 26 (20%) | 128.0 (40.2) | 122.3 (48.9) | 4.8 (43.5) |
| Advanced (III/IV) | 91 (71%) | 154.6 (42.5) | 124.0 (52.8) | 17.2 (42.8) |
|  |  | ***p* = 0.01** | *p* = 0.44 | *p* = 0.21 |
| Response to Therapy |  |  |  |  |
| Complete | 96 (78%) | 147.2 (38.7) | 123.4 (60.7) | 10.4 (45.0) |
| Incomplete | 27 (22%) | 154.4 (57.7) | 120.5 (57.4) | 17.6 (47.1) |
|  |  | *p* = 0.39 | *p* = 0.89 | *p* = 0.48 |
| Surgical Debulking |  |  |  |  |
| Optimal (NED or <1mm) | 98 (81%) | 146.0 (44.1) | 125.2 (59.5) | 9.5 (45.2) |
| Suboptimal (>1mm) | 23 (19%) | 157.5 (45.4) | 113.2 (43.3) | 27.2 (36.0) |
|  |  | *P* = 0.19 | *p* = 0.68 | *p* = 0.05 |
| Survival |  |  |  |  |
| Long-term (>60 months) | 59 (57%) | 140.9 (36.4) | 122.2 (48.6) | 6.9 (40.0) |
| Short-term (≤36 months) | 45 (43%) | 154.6 (44.2) | 121.6 (49.1) | 26.5 (37.2) |
|  |  | ***p* = 0.02** | *p* = 0.83 | ***p* = 0.01** |

NED=No evidence of disease. P-values separately reported for DS and TE were calculated from Wilcoxon rank sum test comparing Cav-1 between clinical subgroups. P-values reported for mean difference were calculated from Student's t-test comparing mean difference between clinical subgroups and for overall test of location (mean=0).
<sup>a</sup> Some variables do not total to 128 due to missing data (age, n=2; stage, n=5; surgical debulking, n=7; survival, n=24).

### 3.4 Expression of Cav-1 using immunofluorescence, Cohort 2

Cav-1 expression in stroma and epithelium regions was quantified using immunofluorescence staining for both benign ovarian and OvCa tissues (Figure 3). Similar to the IHC analysis, the expression of Cav-1 in benign stroma was significantly higher than in the tumor stroma, with median AQUA^®^ scores of 308.9 and 152.6, respectively (p=8.3×10^-13^). However, within tumors, we observed the opposite of IHC analysis and found stromal Cav-1 expression was higher than in the epithelium with median AQUA^®^ scores of 152.6 and 124.0, respectively (p=0.01).

**Figure 3.**
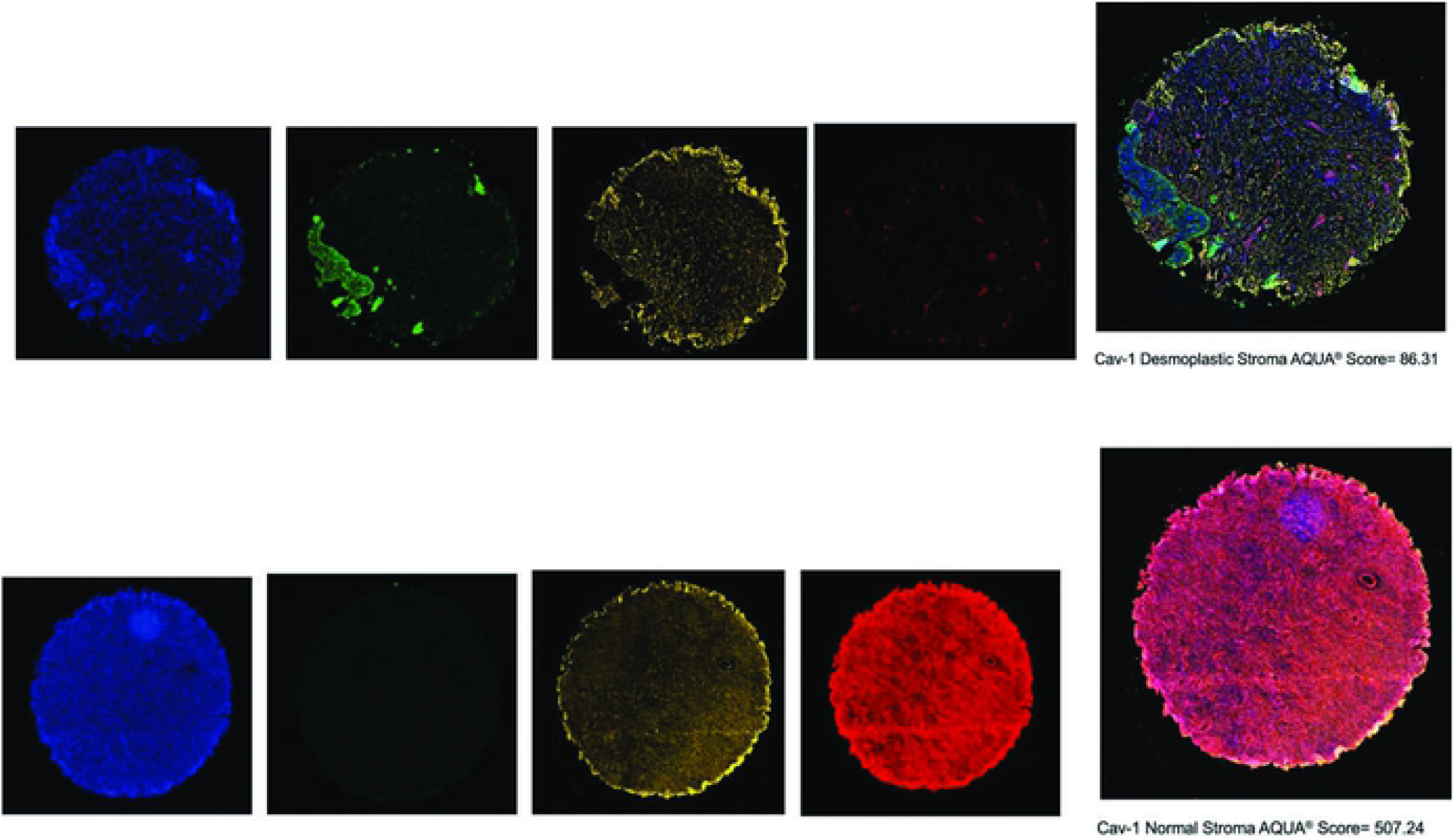
Multiplex immunofluorescent staining of ovarian tissues in Cohort 2. Representative staining for high-grade ovarian serous adenocarcinoma (top) and benign ovarian tissue (bottom) are provided. Compartmental regions were identified based on DAPI, PanCK, and DAPI staining. Tumor (desmoplastic) stroma regions comprised stromal cells (Vimentin+) with a nucleus (DAPI+), and not epithelial (panCK-).

For 11 OvCa tissues quantified by both IF and IHC assays, we observed moderate correlation between methods for Cav-1 expression in stroma (pearson *r*^*2*^= 0.69, p=0.006) whereas there was little correlation for the epithelium (pearson *r*^*2*^=0.006, p=0.98). Notably, 6 out 7 samples that had no stromal Cav-1 expression by IHC (Hscore=0) had >100 AQUA scores (range=102-200, Figure S1a). Among eight samples with moderate IHC expression of Cav-1 in the tumor epithelium (Hscore=3), we observed a wide range of AQUA scores (range=88-257, Figure S1b). Furthermore, stroma expression was only downregulated in 3 samples by IF assay but was downregulated in all samples by IHC (Figure S1c).

Stromal Cav-1 expression was significantly higher for cases over 65 years of age compared to those 65 years and younger (p=0.03) while the expression in the epithelium did not differ by age (p=0.23; Table 2). The high-grade tumors had lower expression of Cav-1 in both the stroma and epithelium compared to low-grade tumors; however, there were only 5 low-grade cases and the difference was not statistically significance (p=0.55 and p=0.14, respectively). Compared to optimally debulked tumors, suboptimal debulked tumors had a larger difference (downregulation) in epithelial Cav-1 (p=0.04). Neither the stroma nor the epithelium expression differed by stage, therapy response, or survival groups.

For patients with low levels (< median) of epithelial Cav-1 expression, the median survival was 57.5 months compared to 50 months for those with high levels (≥ median) (log-rank p=0.83, Figure 4a). For the stroma regions, the median survival was 62.2 months for patients with low levels of Cav-1 expression compared to 46.5 months for those with high levels (log-rank p=0.16, Figure 4b). Cox-proportional hazards modeling of continuous Cav-1 expression with adjustment for age, stage, and response to therapy, found neither epithelial (HR=1.00, 95% CI=0.99-1.01, p=0.97) nor stromal (HR=1.01, 95% CI=0.99-1.02, p=0.32) Cav-1 expression were associated with patient survival.

**Figure 4.**
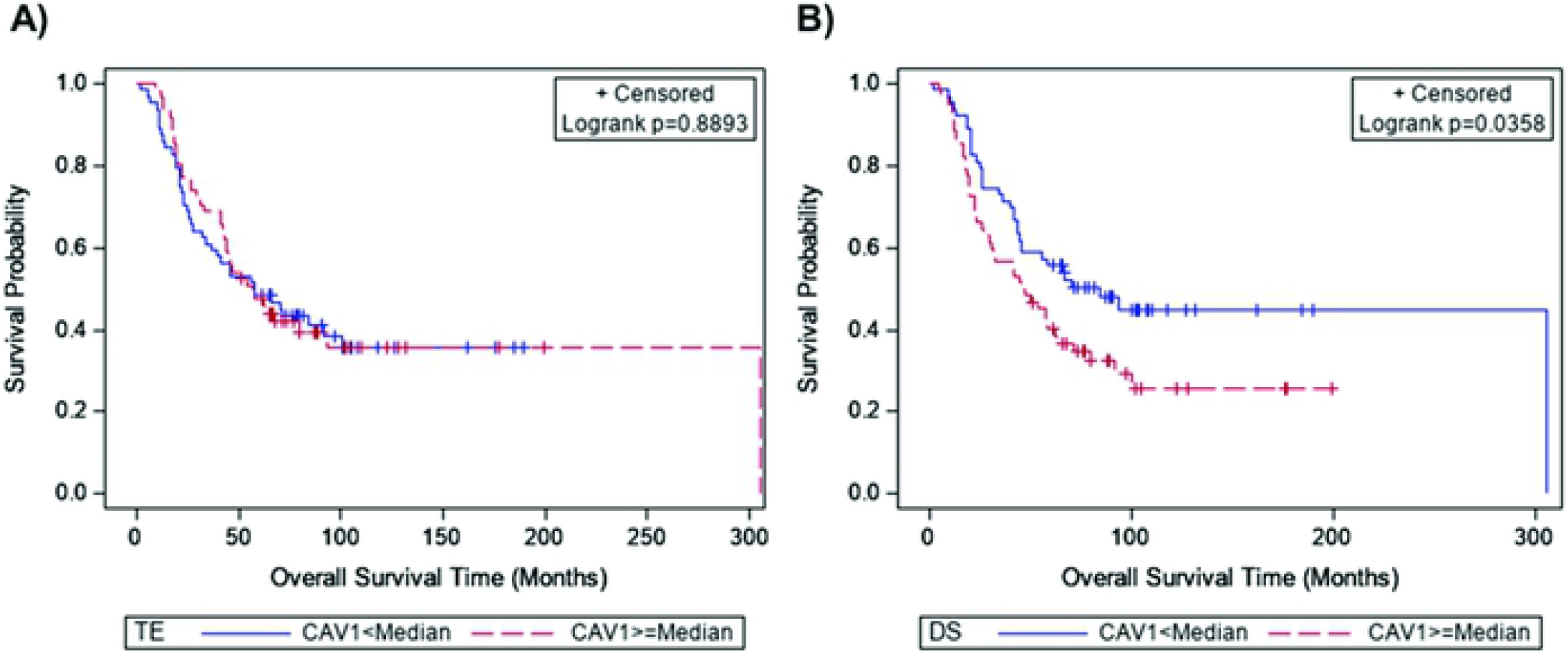
Overall survival by expression of Cav-1 in Cohort 2. Kaplan-Meier (KM) survival curves for Cohort 2 by immunofluorescent expression of Cav-1 in the (A) tumor epithelium and (B) stroma, using the median AQUA^®^ score as the cutoff.

## Discussion

Ovarian cancer remains one of the most lethal gynecological malignancies, with most patients presenting with advanced stage tumors. Unfortunately, current standard treatment regimens are not optimal. Given the emerging importance of the tumor microenvironment with respect to carcinogenesis and chemoresistance, we evaluated the expression of Cav-1, a key potential regulator, specifically exploring its role in ovarian serous adenocarcinoma as a therapeutic and prognostic factor. Our results show that expression of Cav-1 is downregulated in OvCa stroma compared to benign stroma which was detected with both IHC and IF quantification methods. This finding is in-line with an intriguing paradigm of ‘parasitic’ tumor-stromal metabolism called the Reverse Warburg Effect (RWE) [33] which suggests that tumor cells ‘hijack’ CAFs by inducing oxidative stress and downregulating stromal Cav-1 to promote catabolic processes that supply nutrients to tumor cells for anabolic growth and survival thereby promoting tumor progression, metastasis, and resistance to therapy. Cav-1 and other markers of the RWE represent promising targets for new combinations of therapies that work by ‘uncoupling’ tumor cells from CAFs by preventing stromal Cav-1 degradation [34, 35]. Studies in OvCa support the efficacy of these drugs in combination with platinum therapy [34-38] and recent in vitro data shows tumor-stromal metabolic coupling and a loss of stromal Cav-1 in OvCa cell lines [37]. Our findings demonstrate that loss of Cav-1 expression in CAF-enriched stromal cells is a marker of malignant ovarian disease and suggest Cav-1 may be a therapeutic target for OvCa.

The expression of stromal Cav-1 relative to the epithelium varied by assay with IHC showing downregulation in stroma whereas IF showed upregulation. The difference between replicates may be due to tumor heterogeneity or an increased sensitivity and specificity of IF with digital image analysis. In the comparison of biological replicates, IF detected stromal Cav-1 expression in all 7 tissues that displayed no expression with IHC. We also observed a wide-range of epithelium Cav-1 expression detected with IF (88 to 258) for a single IHC Hscore (3). This could result from increased reliability of using objective image analysis versus potentially more subjective human observers. The human eye is more prone to errors when scoring stained slides due to the difference in each individual’s perception of color and intensity and the context in which the human eye views an image, as well as fatigue; whereas, the machine uses individual pixel intensity to score a slide which is an intrinsic property of the image and is not influenced by its surrounding [32]. Given the advancements in digital pathology, future studies should be carried out to promote reproducibility; thus, allowing for improved precision of biomarker measurements in the field.

With the IF assay, we were able to detect differences in Cav-1 expression across clinicopathologic characteristics. We observed higher expression of stromal Cav-1 with age in cohort 2. Additionally, low-grade tumors appeared to have higher expression of Cav-1 than high-grade tumors, although we evaluated only 5 (5%) OvCa tissues given the rare prevalence of low-grade disease at diagnosis (5-10%) [39]. Based on the RWE, one could reasonably extrapolate that as a cancer becomes more aggressive there is more stromal metabolism, and as such, less Cav-1 in the surrounding stroma. However, we also observed a suggested trend (p=0.08) of higher stromal Cav-1 expression among short-term survivors. Perhaps, in advanced stage tumors that are associated with shorter survival times, changing metabolic needs [40] and migration of aggressive tumor cells require less nutrients from the surrounding stroma, while in the early stage tumors which are associated with longer survival times, the tumor cells remain locally and deplete the surrounding stomal Cav-1. We also evaluated the association of Cav-1 expression with response to standard first-line platinum-based chemotherapy but did not detect a difference. Taken together, these findings suggest a limited clinical utility for stromal Cav-1 expression as a biomarker for predicting chemotherapy response.

Overall, the Cav-1 expression levels in the tumor microenvironment are quantifiable via digital image analysis and could potentially serve as a diagnostic biomarker assay. Given that only malignant (or benign) cases were assessed, future studies should be undertaken to explore the utility of Cav-1 in more diagnostically challenging, “borderline” ovarian neoplasms. In-line with previous studies [12-14, 26], the statistically significantly higher expression of Cav-1 in the benign stroma versus tumor stroma demonstrates that Cav-1 may play an important role in carcinogenesis and further studies may provide insight into the evolutionary biology of OvCa. Although the utility in predicting response to standard platinum-based chemotherapy was limited, Cav-1 could prove to be an interesting therapeutic target based on its loss in malignant ovarian tissue. Studies are needed to glean mechanistic insights into how reduced levels of Cav-1 in the stroma may promote transformation to ovarian carcinoma.

Our findings demonstrate that loss of desmoplastic stromal Cav-1expression may be associated with malignant ovarian serous carcinomas. Studies are needed to replicate these findings and explore mechanisms that can be targeted to reduce progression to malignancy.

## Data Accessibility

The data from this manuscript is not in a public database or repository.

## Author Contribution

Conceptualization: JBP, AM

Data Curation: DS-V, BR, DM, SM, AH, AL, DT-C, JBP

Formal analysis: DS-V, BR, DM, SM, AH, AL, DT-C, JBP

Funding acquisition: JBP, JML

Investigation: DS-V, BR, DM, SM, AH, AL, RW, SA, DT-C, AM, JBP

Methodology: JBP, AM, DS-V, BR

Project Administration: JBP, DS-V, BR

Resources: JBP, JML

Software: DS-V

Supervision: JBP, AM, JML

Validation: DS-V, BR, DM, SM, AH, AL, DT-C, JBP

Visualization: DS-V

Writing-original draft preparation: DS-V, BR, JBP, AM

Writing-review and editing: DS-V, BR, DM, SM, AH, AL, RW, SA, DT-C, AM, JBP

## Acknowledgements

The authors wish to thank the study participants for donating tissue and data for this project.

## SUPPORTING INFORMATION

**Supplemental Figure S1**. Comparison of Cav-1 expression quantified by immunofluorescence and immunohistochemistry.

**Supplementary IF staining:**

The first step of the protocol consists of primary antibody incubation using a vimentin mouse monoclonal antibody (Abcam Biotechnology lot: ab8069, Cambridge, UK), to visualize the fibroblasts in the stroma, at a 1:80000 dilution for 30 minutes. This is followed by the labeled polymer incubation step using a mouse envision reagent (Dako ref: K4007) for 30 minutes and a substrate, TSA-Cy3 (PerkinElmer ref: NEL744B001KT), incubated at a 1:50 dilution for 5 minutes. After the first round of IF staining the slides were washed in a 0.1% azide solution and the staining process was repeated this time using a CAV-1 rabbit polyclonal antibody (Santa Cruz lot: G0314) at a 1:5000 dilution and a guinea pig pancytokeratin tumor masking antibody (Acris lot: 411101, Rockville, MA) at a 1:100 dilution for 30 minutes. This was followed by the labeled polymer incubation step using a rabbit envision reagent (Dako ref: K4011) and a secondary antibody, Alexa488 guinea pig (Abcam, ab150185), to visualize the tumor masking antibody, at a 1:200 dilution for 30 minutes and a substrate TSA-Cy5 (Perkin Elmer ref: NEL745B001KT) incubated at a 1:50 dilution for 5 minutes. The slides were cover-slipped with ProLong Gold antifade reagent with 4’,6-diamidino-2-phenylindole (DAPI) mounting media to visualize the nuclei.

